# Pan-Cancer Driver Mutation Signatures Define a Molecular Taxonomy of Tumors

**DOI:** 10.64898/2026.06.17.732825

**Authors:** Xiaoyun Huang, Bowang Chen, Xiaoqing Huang, Martin C.S. Wong

**Author notes:** Correspondence: Xiaoyun Huang and Martin C.S. Wong.

## Abstract

Cancers with similar histology often exhibit divergent clinical behavior, reflecting molecular heterogeneity not captured by current classification systems. Although driver mutations are central to tumorigenesis, their broader systems-level consequences have not been systematically leveraged. We integrated genomic and transcriptomic data across cancers to define driver mutation signatures (DMS), coordinated transcriptional programs associated with cancer driver mutations. From 121 candidate drivers, we derived 90 robust signatures and quantified their activity in individual tumors using mutation signature scores (MSS). DMS analysis revealed a hierarchical organization of tumors into molecular subgroups that transcended tissue boundaries while preserving driver-associated features. Continuous MSS profiles further defined high-resolution molecular fingerprints for individual tumors. DMS provides a quantitative framework for tumor classification and patient stratification and links driver-associated programs to potential therapeutic vulnerabilities. Together, these findings establish a pan-cancer molecular taxonomy that bridges genotype and phenotype and may inform precision oncology.

## Introduction

Cancers that appear histologically similar often exhibit profoundly different behaviours, diverging in their evolutionary trajectories, therapeutic vulnerabilities and clinical outcomes. This variability reflects not only the accumulation of somatic mutations, but also how specific driver mutations reshape cellular programs and tumor ecosystems (*1–10*) . Although individual driver genes such as TP53, KRAS, and EGFR have been extensively studied, it remains unclear whether tumors can be systematically organized into a unifying molecular taxonomy based on the combinatorial impact of driver mutations across cancer types.

Large-scale sequencing efforts have transformed our ability to characterize genomic alterations at unprecedented resolution. Yet, a central paradox persists: despite the abundance of genomic data, clinically actionable stratification remains limited. Canonical classification systems, largely defined by tissue of origin and histopathology, fail to reflect the molecular logic that governs tumor behavior (*11*). Even within molecular frameworks, most approaches focus on single mutations or mutational burden , overlooking the broader transcriptional and functional consequences of driver events (*12–17*). As a result, key questions remain unresolved: Do driver mutations impose consistent, system-level effects on tumor biology? And can these effects be leveraged to define cross-cancer subtypes that transcend tissue boundaries?

Emerging evidence suggests that driver mutations do not act in isolation, but rather generate coordinated transcriptional programs, referred to here as driver mutation signatures, that capture their downstream biological impact. These signatures may encode information about proliferative capacity, immune evasion (*18*), metabolic rewiring (*19*), and other hallmarks of cancer (*20–23*). However, their potential to organize tumors into a coherent, pan-cancer framework has not been systematically explored. In particular, it is unknown whether such signatures can reveal latent structure within the cancer landscape, enabling a classification scheme that is both biologically meaningful and clinically informative.

Here, we address this gap by constructing a comprehensive atlas of driver mutation signatures across cancers and using these signatures to define a molecular taxonomy of tumors. By integrating genomic and transcriptomic data at scale, we seek to move beyond mutation-centric views toward a systems-level understanding of how driver events shape tumor identity. This framework provides a foundation for reclassifying cancers based on shared molecular programs, with implications for patient stratification, therapeutic targeting, and the broader principles governing tumor heterogeneity.

## Results

### Driver Mutation Signatures Encode System-Level Programs and Define Tumor Subtypes

We present an integrated pipeline for identifying Molecular Subtypes of Tumors based on driver mutation signature (DMS) quantification (**fig. S1**). We focused on a list of 121 statistically characterized pan-cancer driver genes (*24*). First, differential expression analysis was performed to identify the signature genes for each cancer driver gene, resulting in robust signatures for 90 cancer driver genes. To further characterize the biological programs captured by cancer driver gene mutation, we aggregated all signature gene sets derived from 90 cancer drivers, yielding a total of **2,904** unique genes. These genes were subjected to functional enrichment analysis using Metascape (*25*), which revealed a highly structured network of interconnected biological processes (**fig. S2**). Besides canonical pathways such as cell cycle and pathways in cancer, driver mutations actively modify the immune system and extracellular matrix. Importantly, the network of enriched pathways revealed clear modular organization, with distinct clusters corresponding to immune, stromal, and proliferative processes, yet interconnected through shared gene memberships. This indicates that mutation-derived transcriptional signatures are not isolated features but instead reflect coordinated, system-level biological programs.

The pipeline further included downstream analysis to identify key driver mutation signatures associated with each subtype, providing a comprehensive framework for classifying tumor samples based on their driver mutation signatures. We estimated the number of clusters using elbow method and silhouette method (**fig. S3**). As a result, we obtained a DMS based cancer subtypes (S1-S8) (**fig. 1A**). We then interrogated the diversity of DMS subtypes within TCGA projects using Gini index and Shannon entropy, revealing that ACC and COAD are among the most diversified cancer types (**fig. 1B**). We visualized the top five cancer driver mutation signatures for each DMS subtype, characterized by distinct patterns (**fig. 1C**). TP53 and RB1 mutation signatures co-exist among the top five for S1, S3, S4 and S8. Three RAS genes are among top cancer driver signatures. High KRAS mutation signature was a hallmark for S1, S4 and S6; while high NRAS mutation signature was observed in S3 and S8. Besides S4 and S7, all subtypes have HRAS mutation signature to some extent. Interestingly, S7 seemed to be RAS independent. Alternative splicing related gene SF3B1 mutation signature is most obvious in S8, which consists of blood cancers. We furthered investigated the top genes expressed in each DMS subtype, revealing crystal-clear distinct top markers for each subtype (**fig. 1D**). S1 is characterized by upregulation of KRT genes, including KRT6A, KRT14, KRT5, KRT17, KRT16, KRT13 and KRT6B. S2 is enriched for neuroendocrine related genes CHGA and CHGB.

**Fig. 1.**
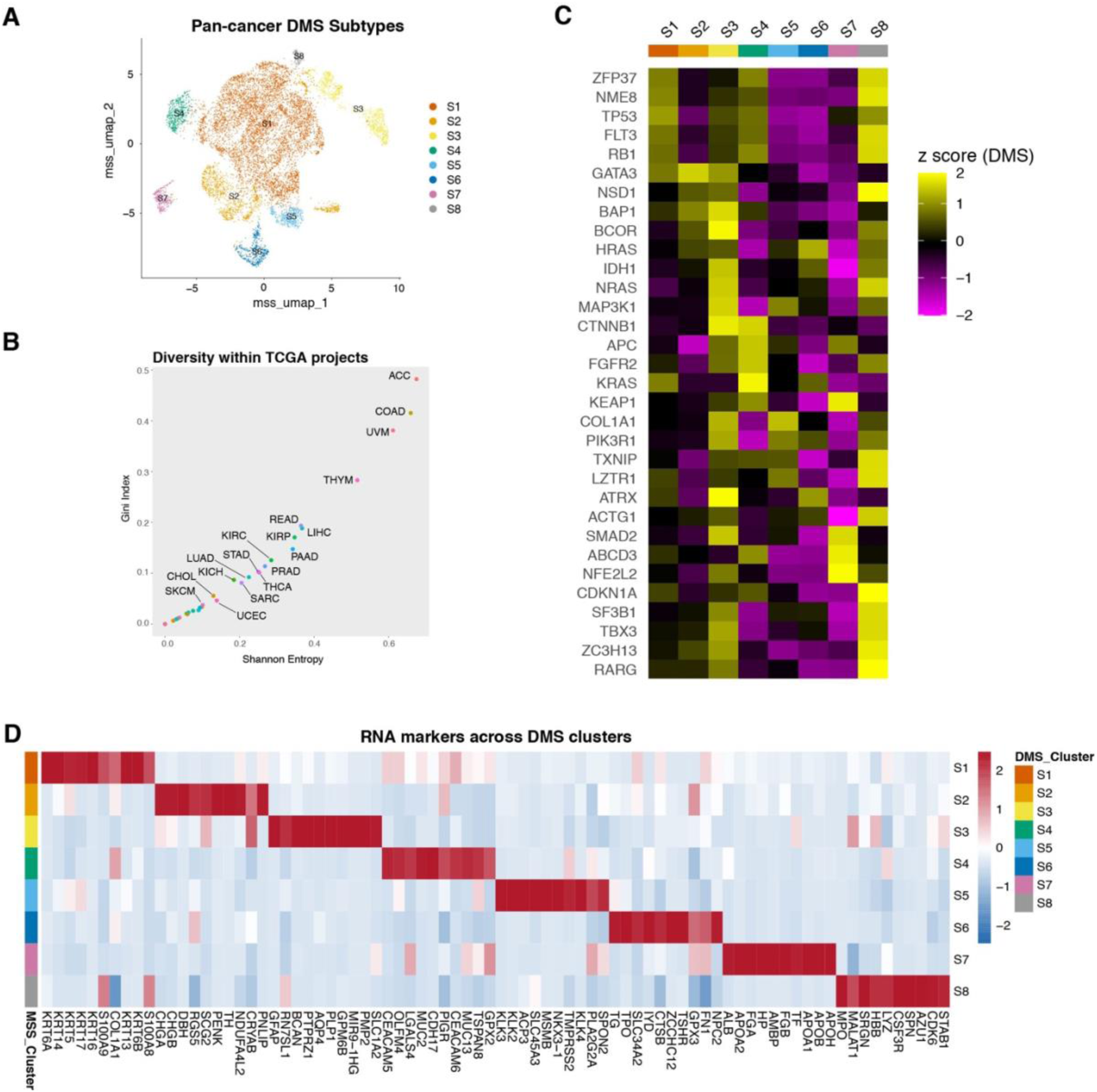
Pan-cancer landscape of driver mutation signature (DMS) subtypes and associated molecular features. (**A**) Uniform Manifold Approximation and Projection (UMAP) embedding of TCGA tumor samples based on mutation signature scores (MSS) derived from 90 driver mutation signatures. Each point represents an individual tumor sample, colored by DMS-defined subtype (S1–S8). (**B**) Scatter plot showing diversity of DMS subtype composition within each TCGA cancer type. Each point represents a cancer type, positioned by Shannon entropy (x-axis) and Gini index (y-axis), reflecting subtype diversity and inequality, respectively. (**C**) Heatmap displaying the average standardized mutation signature scores (Z-score) for representative driver genes across DMS subtypes (S1–S8). Rows correspond to driver genes, and columns correspond to subtypes. (**D**) Heatmap of RNA expression markers across DMS subtypes. Each row represents a subtype (S1–S8), and each column represents a gene. Expression values are scaled (Z-score) across samples.

### DMS subtyping stratifies patient survival and clinical risk across cancers

To evaluate the clinical relevance of DMS-defined subtypes, we performed Kaplan–Meier survival analysis across TCGA cohorts using the well annotated survival data (*26*). Patients stratified by the eight DMS clusters exhibited markedly distinct overall survival (OS) outcomes (log-rank test, p < 0.0001; **fig. 2A**), demonstrating strong prognostic separation. Among all subtypes, S8 was associated with the poorest survival, characterized by a rapid decline in survival probability, whereas S5 showed the most favorable prognosis with sustained long-term survival. Within the more prevalent subtypes, S2 displayed relatively favorable outcomes, while S3 was associated with inferior survival, indicating heterogeneity even among dominant groups.

**Fig. 2.**
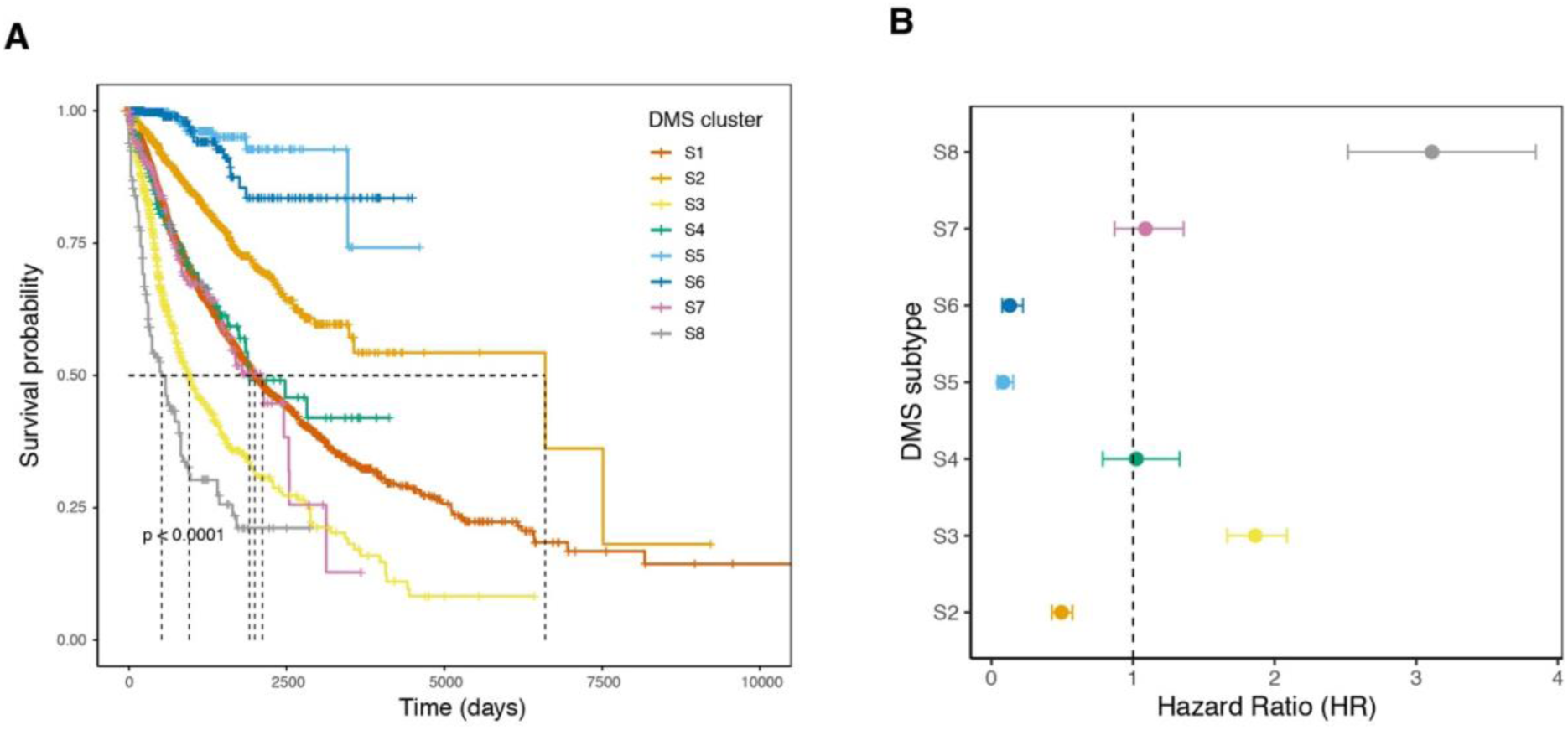
Clinical relevance of DMS subtypes across pan-cancer cohorts. (**A**) Kaplan–Meier overall survival curves for TCGA patients stratified by DMS-defined subtypes (S1–S8). Each curve represents a subtype, with tick marks indicating censored observations. The dashed horizontal line marks 50% survival probability, and vertical dashed lines indicate median survival times for each subtype. Significant differences in survival outcomes are observed across subtypes (log-rank test, p < 0.0001), demonstrating the prognostic value of DMS-based classification. (**B**) Forest plot showing hazard ratios (HRs) and 95% confidence intervals for each DMS subtype derived from Cox proportional hazards regression. The dashed vertical line indicates HR = 1 (reference).

To quantify subtype-specific risk, we further performed Cox proportional hazards analysis (**fig. 2B**). Consistent with the survival curves, S8 exhibited the highest hazard ratio (HR), indicating significantly increased mortality risk, whereas S5 and S6 were associated with reduced risk (HR < 1). Other subtypes showed intermediate or near-neutral effects, with variability in confidence intervals reflecting subtype-specific heterogeneity. These results establish DMS subtyping as a robust prognostic framework that captures clinically meaningful differences in survival risk across cancers.

### Association between driver mutation signatures and tumor mutational burden

To investigate the relationship between driver mutation signatures and genomic instability, we systematically correlated mutation signature scores (MSS) with tumor mutational burden (TMB) across TCGA samples. Globally, most driver-associated signatures exhibited positive correlations with TMB, indicating that mutation-driven transcriptional programs are broadly linked to increased mutational load (**fig. 3A**). Among these, TP53 emerged as one of the top-ranked drivers, alongside genes such as ZFP37 and ZNF624, showing strong positive associations with TMB.

**Fig. 3.**
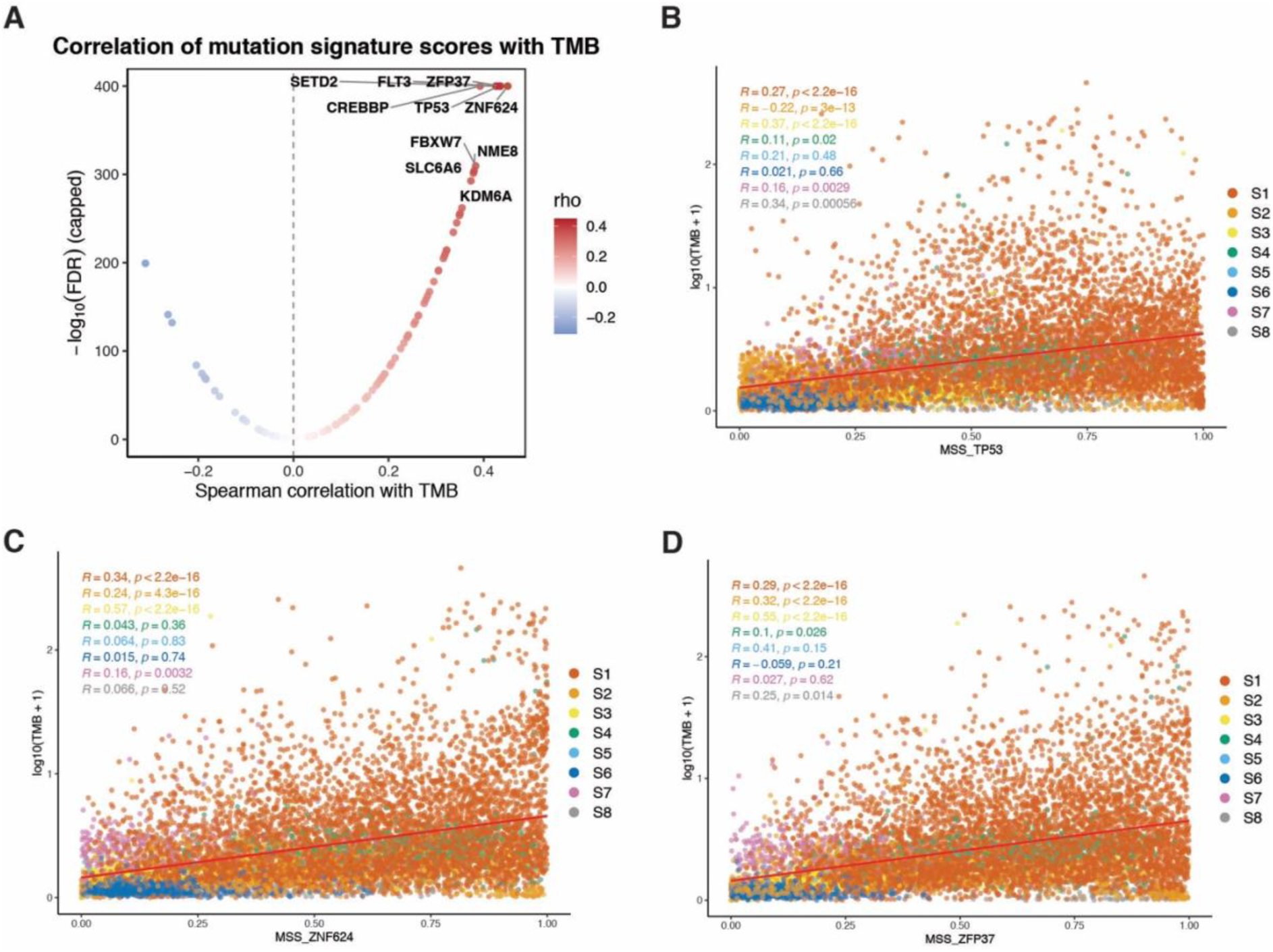
Association between driver mutation signature scores and tumor mutational burden (TMB). (**A**) Each point represents a driver mutation signature. The x-axis shows the Spearman correlation coefficient (ρ) between mutation signature score (MSS) and TMB across TCGA samples, and the y-axis shows the statistical significance (−log10 FDR, capped). Positive correlations dominate, indicating that many driver-associated transcriptional programs are linked to increased mutational burden. Selected representative drivers are annotated. (**B–D**) Representative associations between MSS and TMB at the sample level. Scatter plots showing the relationship between TMB (log10-transformed) and MSS for representative driver signatures (B, TP53; C, ZNF624; D, ZFP37). Each point represents a tumor sample, colored by DMS subtype (S1–S8). Red lines indicate linear regression fits. Subtype-specific correlation coefficients (R) and significance levels are shown.

At the sample level, MSS for TP53 displayed a consistent positive correlation with TMB across tumors (**fig. 3B**), supporting a central role of TP53-associated transcriptional programs in shaping genomic instability. Similar patterns were observed for additional high-ranking drivers, including ZNF624 and ZFP37 (**fig. 3C–D**), although the strength of association varied across DMS subtypes. Notably, subtype-specific analyses revealed heterogeneous correlation patterns, suggesting that the coupling between mutational burden and transcriptional output is modulated by the broader molecular context. Those results demonstrate that DMS captures biologically meaningful signals linked to tumor mutational processes, with TP53-associated signatures representing a key axis connecting genomic instability and transcriptional reprogramming in cancer.

### Cellular Composition Across DMS Subtypes and Immune Subtypes

To explore the relationship between DMS subtypes and cellular composition, we performed xCell-based deconvolution (*27*) to estimate cell type enrichment scores for each sample in the TCGA dataset (**fig. S4A**). Notably, DMS clusters exhibit distinct cellular profiles compared to other clusters, with significant differences in the abundance of various cell types, including immune and stromal cells. We further interrogated the importance of each inferred cell types in DMS subtypes. We ranked the cell types in distinguishing the DMS clusters based on feature importance scores derived from a Random Forest model (**fig. S4B**). This analysis revealed that Keratinocytes and Megakaryocytes were among the most important features, highlighting the strong influence of these cell types in the overall DMS subtype classification.

We further investigated the abundance of major cell types (immune cells, stromal cells, endothelial, and epithelial cells) across the DMS clusters (**fig. S4C**). The distribution of xCell scores for these cell types demonstrates that specific DMS clusters are enriched in distinct cell populations. For instance, S1 shows high scores for Epithelial cells, whereas S2 exhibits strong enrichment for Endothelial cells.

We next explored the relationship between DMS subtypes and the previously proposed immune subtypes (Immune_Subtype) (*28*) through a Sankey diagram, demonstrating how DMS clusters are distributed across the immune subtypes (**fig. S4D**). Despite the distinct cellular composition of each DMS subtype, the immune microenvironment (as represented by Immune_Subtype) does not fully align with the DMS subtypes. All DMS subtypes consists of several immune subtypes (C1-C6), suggesting that immune composition alone does not fully capture the heterogeneity of DMS subtypes. These findings underscore that DMS subtypes are driven by a complex combination of cellular composition, including immune and stromal cells, and are distinct from cancer subtypes based solely on immune composition. Thus, DMS subtyping provides a deeper, more nuanced stratification of cancer that incorporates both mutational profiles and the tumor microenvironment, highlighting the potential of DMS as a clinically relevant marker for tumor heterogeneity and immune interactions.

### Fine Subclustering of DMS S1 Subtype Reveals Clinically Relevant Molecular Heterogeneity

The DMS S1 cluster, which contains the largest proportion of samples, exhibits substantial intra-cluster heterogeneity, suggesting the need for finer stratification of this group (**fig. S5**). We aimed to investigate the heterogeneity within this dominant DMS subtype (S1) and its clinical and molecular implications. By performing a finer subclustering analysis on the S1 samples, we identified eight distinct subclusters (S1.1–S1.8) (**fig. 4A**). These subclusters represent molecularly diverse subtypes within the DMS S1 cluster, offering valuable insights into their clinical significance and underlying biological processes.

**Fig. 4.**
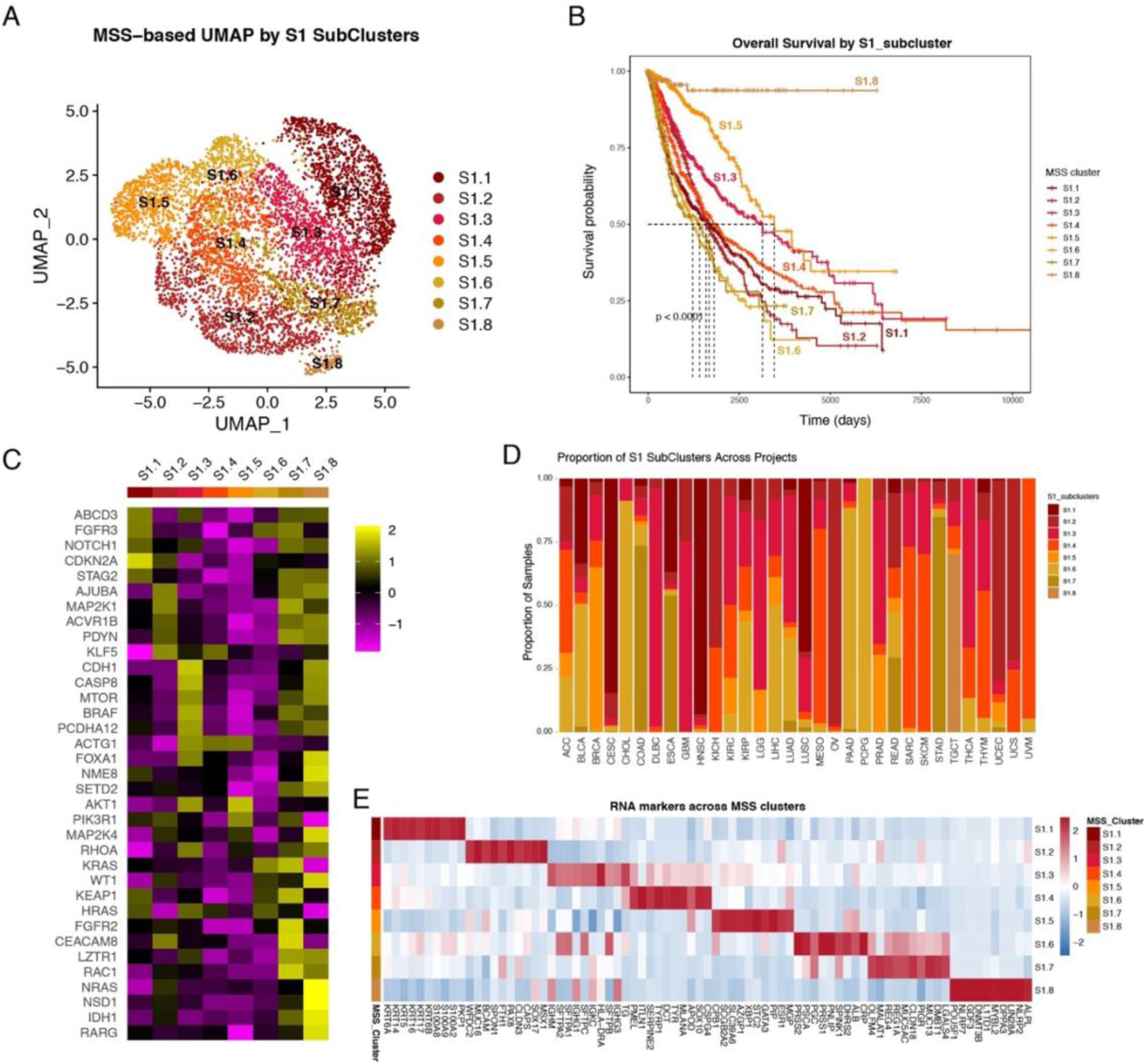
Refinement of S1 into distinct subclusters reveals additional molecular and clinical heterogeneity. (**A**) UMAP embedding of tumor samples within the S1 cluster using mutation signature scores (MSS), revealing eight distinct subclusters (S1.1–S1.8). Each point represents an individual sample, colored by subcluster assignment, highlighting substantial intra-cluster heterogeneity within S1. (**B**) Kaplan–Meier overall survival curves for patients stratified by S1 subclusters (S1.1–S1.8). Tick marks indicate censored observations. The dashed horizontal line denotes 50% survival probability, and vertical dashed lines indicate median survival times. Significant survival differences are observed across subclusters (log-rank test, p < 0.0001), demonstrating the clinical relevance of finer stratification within S1. (**C**) Heatmap showing standardized mutation signature scores (Z-score) of representative driver genes across S1 subclusters. Rows represent driver genes and columns represent subclusters. (**D**) Stacked bar plot showing the proportion of S1 subclusters across TCGA cancer types. Each bar represents a cancer type, and colors indicate the relative contribution of each subcluster, revealing cancer-type-specific enrichment patterns. (**E**) Heatmap of RNA expression markers across S1 subclusters. Rows represent subclusters and columns represent genes. Expression values are scaled (Z-score) across samples.

We next assessed the overall survival (OS) across these subclusters (**fig. 4B**). Kaplan-Meier survival analysis revealed significant differences in survival outcomes between the subclusters (p < 0.0001), highlighting the clinical heterogeneity within the S1 subtype. For example, S1.8, S1.5, and S1.3 subclusters exhibited notably better survival compared to other subclusters, indicating that subclustering within the MSS S1 subtype can stratify patients based on their survival probabilities.

To better understand the biological underpinnings of the identified subclusters, we examined the top driver mutation signatures across these subtypes (**fig. 4C**). The heatmap of top driver mutation signatures revealed that each S1 subcluster is associated with a distinct set of oncogenic drivers. For instance, S1.7 exhibited high mutation signature of FGFR2. Oncogenic mutation of FGFR2 has been associated with drug resistance. Targeting FGFR2 with inhibitors might improve the dismal prognosis for this DMS subtype. These findings indicate that the molecular landscape of the S1 subtype is shaped by a diverse set of tumorigenic pathways, further distinguishing the subclusters from one another.

The prevalence of these subclusters across different cancer types was also investigated (**fig. 4D**). This distribution emphasizes the broad applicability of these S1 subclusters in various cancers and underscores the potential utility of finer subtyping for patient stratification in clinical settings.

Given recent evidence implicating uPAR (encoded by PLAUR) as a key feature of tumors harboring TP53 and RAS pathway alterations (*29*), we examined the relationship between PLAUR expression and mutation signature scores (MSS) for TP53 and KRAS. Across TCGA samples, PLAUR expression was elevated in tumors with high TP53 and KRAS MSS, with the highest expression observed in the upper-right region of the MSS_TP53–MSS_KRAS space (**fig. S6A**). This pattern indicates a cooperative association between TP53- and KRAS-driven transcriptional programs and PLAUR upregulation. Consistent with this observation, stratification within the dominant S1 lineage revealed marked heterogeneity of PLAUR expression across subclusters, with specific subgroups (e.g., S1.6 and S1.4) exhibiting higher expression levels (**fig. S6B**).

### A hierarchical landscape of driver mutation signatures reveals multi-dimensional tumor heterogeneity

To capture both global and fine-grained transcriptional variation, we implemented a two-layer hierarchical DMS framework, in which broad subtypes (S1–S8) are complemented by finer stratification within the S1 lineage (**fig. 5**). Rather than representing a uniform group, S1 decomposes into multiple transcriptionally distinct states, revealing that a substantial fraction of tumors share overarching driver-associated programs while diverging along more subtle regulatory axes.

**Fig 5.**
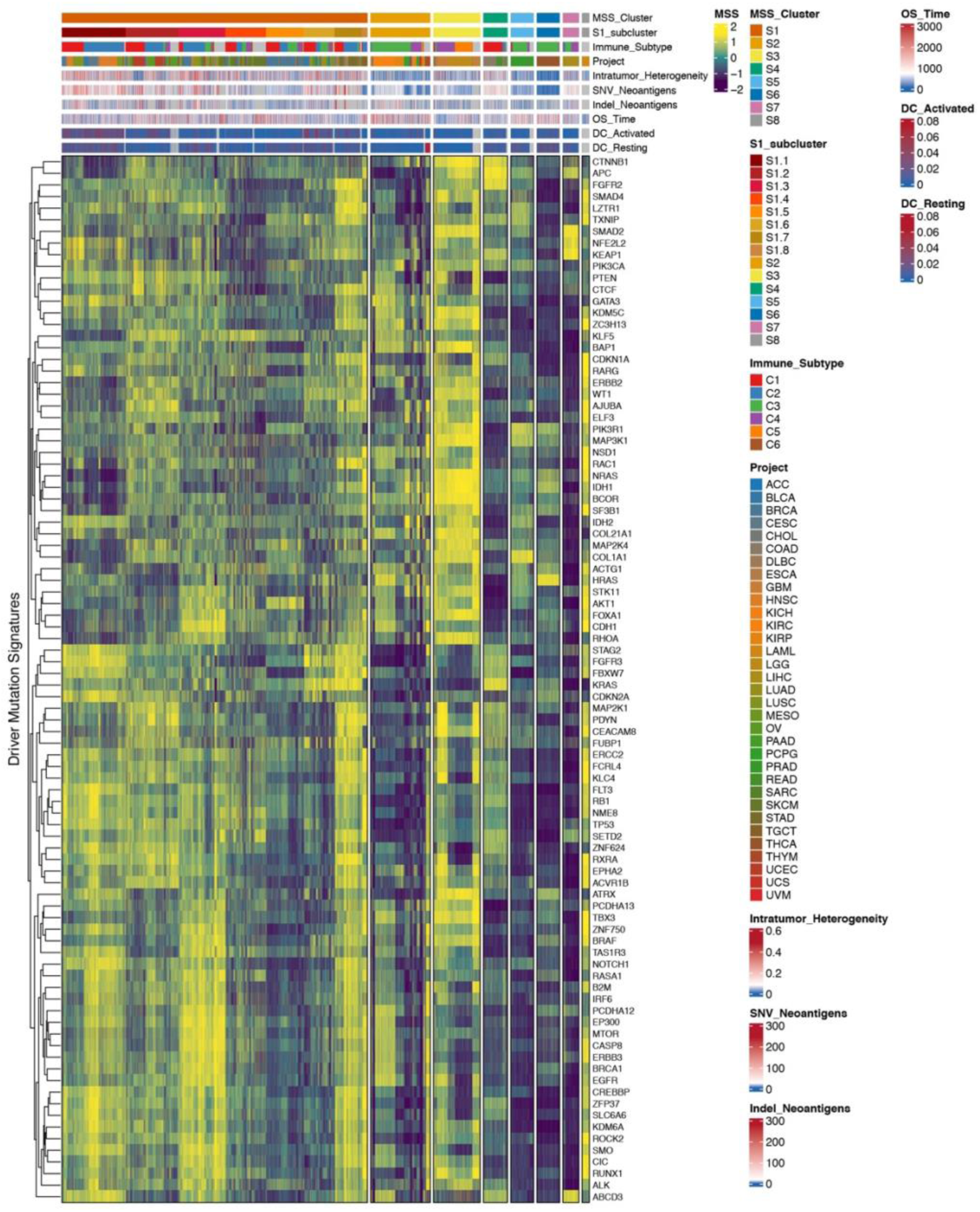
Integrated landscape of driver mutation signatures and multi-dimensional tumor features across pan-cancer cohorts. Comprehensive heatmap summarizing mutation signature scores (MSS) alongside clinical, genomic, and immunological features across TCGA tumor samples. Columns represent individual samples and rows represent driver mutation signatures, hierarchically clustered based on MSS similarity. MSS values are scaled (Z-score) to highlight relative activation patterns. Top annotation tracks display DMS cluster (S1–S8), S1 subcluster classification (S1.1–S1.8), immune subtype (C1–C6), and cancer type (TCGA projects). Additional quantitative tracks include intratumor heterogeneity, single-nucleotide variant (SNV) neoantigen load, insertion/deletion (indel) neoantigen load, overall survival time, and dendritic cell (DC) activation and resting scores.

Integration with additional biological layers shows that this hierarchy is not purely transcriptomic. Subtype-specific variation in intratumor heterogeneity, SNV/indel neoantigen burden, and dendritic cell activity aligns with MSS patterns, demonstrating that DMS captures both driver-induced transcriptional programs and broader genomic instability and antigenicity. Tumors with elevated mutation-associated transcriptional activity tend to have higher predicted neoantigen loads, suggesting a functional link between driver programs and immunogenic potential.

Immune composition is partially concordant with MSS subtypes, yet not fully overlapping, indicating that MSS captures tumor-intrinsic programs distinct from conventional immune classifications. Notably, activated and resting dendritic cell signatures map to specific MSS regions, suggesting mutation-driven transcriptional programs influence antigen presentation and immune priming.

Collectively, these results establish that DMS-based subtyping integrates driver-associated transcriptional programs, genomic instability, immune context, and clinical behavior, providing a multi-dimensional framework for understanding tumor heterogeneity and potential therapeutic vulnerabilities.

### Validation of MSS-based subtype assignment using single-cell RNA-seq projection

To evaluate the generalizability of the mutation signature scoring framework beyond bulk transcriptomic data, we applied our reference-anchored projection strategy to independent single-cell RNA-seq datasets (*30–32*). For each dataset, pseudo-bulk expression profiles were generated by aggregating gene expression across cells within individual samples, followed by projection into the TCGA-derived MSS space using gene-level normalization parameters and driver-specific signature definitions established from the reference cohort.

Across all evaluated datasets, MSS profiles derived from scRNA-seq pseudo-bulk data demonstrated strong concordance with TCGA-derived subtype structures (**fig. S7**). Projection of query samples onto subtype centroids revealed clear subtype preferences, with the majority of samples exhibiting highest correlation to a single DMS subtype, indicating robust preservation of the underlying transcriptional programs associated with driver mutation signatures. Consistent results were obtained using both centroid-based correlation and k-nearest neighbor (kNN) approaches, with high agreement observed for the majority of samples, supporting the stability of subtype assignment across independent methodologies.

### Subtype-specific Drug-driver Interaction Landscapes Reveal Distinct Therapeutic Vulnerabilities

To translate mutation signature–derived driver activity into potential therapeutic insights, we systematically interrogated drug–gene interaction networks using the top-ranked driver signatures for each DMS subtype. By integrating subtype-specific driver prioritization with curated drug–target interaction data, we constructed subtype-resolved therapeutic landscapes that reveal both shared and distinct vulnerabilities across tumors.

Across the eight DMS subtypes (S1–S8), several recurrent oncogenic hubs emerged as highly druggable nodes, including receptor tyrosine kinases and signaling pathway regulators (**fig. S8**). Notably, fibroblast growth factor receptor (FGFR) family members (FGFR2/3), RAS pathway components (KRAS, HRAS), and PI3K pathway regulators (PIK3R1, PIK3CA) were repeatedly identified across multiple subtypes, suggesting convergent targeting opportunities. For example, S1 and S4 subtypes were enriched for FGFR- and KRAS-associated signatures, corresponding to available inhibitors such as erdafitinib, sotorasib, and adagrasib, indicating that these tumors may be broadly susceptible to targeted therapies against receptor tyrosine kinase and MAPK signaling axes .

Despite these shared features, each subtype also exhibited distinct drug–driver interaction profiles. S3 and S6 subtypes were characterized by enrichment of PI3K pathway and chromatin-associated drivers, with corresponding interactions involving PI3K inhibitors (e.g., panulisib) and IDH1-targeted agents, reflecting metabolic and epigenetic dependencies. In contrast, S7 displayed a unique enrichment of oxidative stress–related drivers (KEAP1/NFE2L2 axis) and ERBB3 signaling, suggesting sensitivity to redox-modulating agents and HER-family targeted therapies. S8, which showed relatively distinct transcriptional signatures, was associated with retinoic acid receptor signaling (RARG), highlighting potential responsiveness to retinoid-based therapies.

We next extended this analysis to the finer subclusters within the dominant S1 subtype, which was further stratified into eight subclusters (S1.1–S1.8). This higher-resolution analysis revealed substantial diversification of therapeutic vulnerabilities within S1 (**fig. S9**). While certain pathways remained recurrent—such as FGFR, KRAS, and PI3K signaling—individual subclusters exhibited distinct druggable hubs. For instance, S1.1 and S1.6 were enriched for FGFR3- and FLT3-associated signatures, whereas S1.3 showed strong EGFR/ERBB3 and mTOR pathway activity, and S1.5 was dominated by AKT1 and PI3K signaling. Other subclusters displayed more specialized dependencies, including NOTCH1 signaling in S1.1, MAP2K1 signaling in S1.2 and S1.7, and TP53-associated programs in S1.4 and S1.8 .

Importantly, this subcluster-level analysis revealed that even within a single dominant DMS subtype, tumors may engage distinct oncogenic circuits that map to different therapeutic strategies. For example, while KRAS-targeted therapies may be broadly relevant across several S1 subclusters, others may preferentially respond to FGFR inhibitors, MEK inhibitors, or PI3K/AKT pathway inhibitors, underscoring the need for refined stratification beyond coarse subtype definitions.

Taken together, these results demonstrate that DMS-derived tumor classification can be leveraged to infer subtype-specific drug–target interaction networks, providing a functional bridge between transcriptomic states and therapeutic opportunities. This framework highlights both shared oncogenic dependencies across tumor types and fine-grained heterogeneity within subtypes, supporting the potential of DMS-based stratification to guide precision oncology.

### Extrapolation of Top Cancer Driver Mutation Signatures from RNA-seq Data for Individual Patients

The DMS scores not only enabled tumor classification, but also provide a cancer driver mutation fingerprint for each individual patient. Given individual patient with only RNA-seq data, the top cancer driver mutation signatures can be extrapolated to guide personalized cancer medicine. We visualize an example of how mutation signature scores (DMS) can be derived for individual cancer patients using only their RNA-seq data. The circular heatmap illustrates the DMS scores for a patient (TCGA-A8-A09D) across 90 known cancer driver mutations (**fig. 6A**), revealing *NSD1*, *FUBP1*, and *PCDHA13* as the top three driver mutation signatures. This visualization highlights that, even without access to somatic mutation data, the RNA expression profile of a tumor can provide valuable insights into the driver mutations that may be implicated in its pathology.

**Fig. 6.**
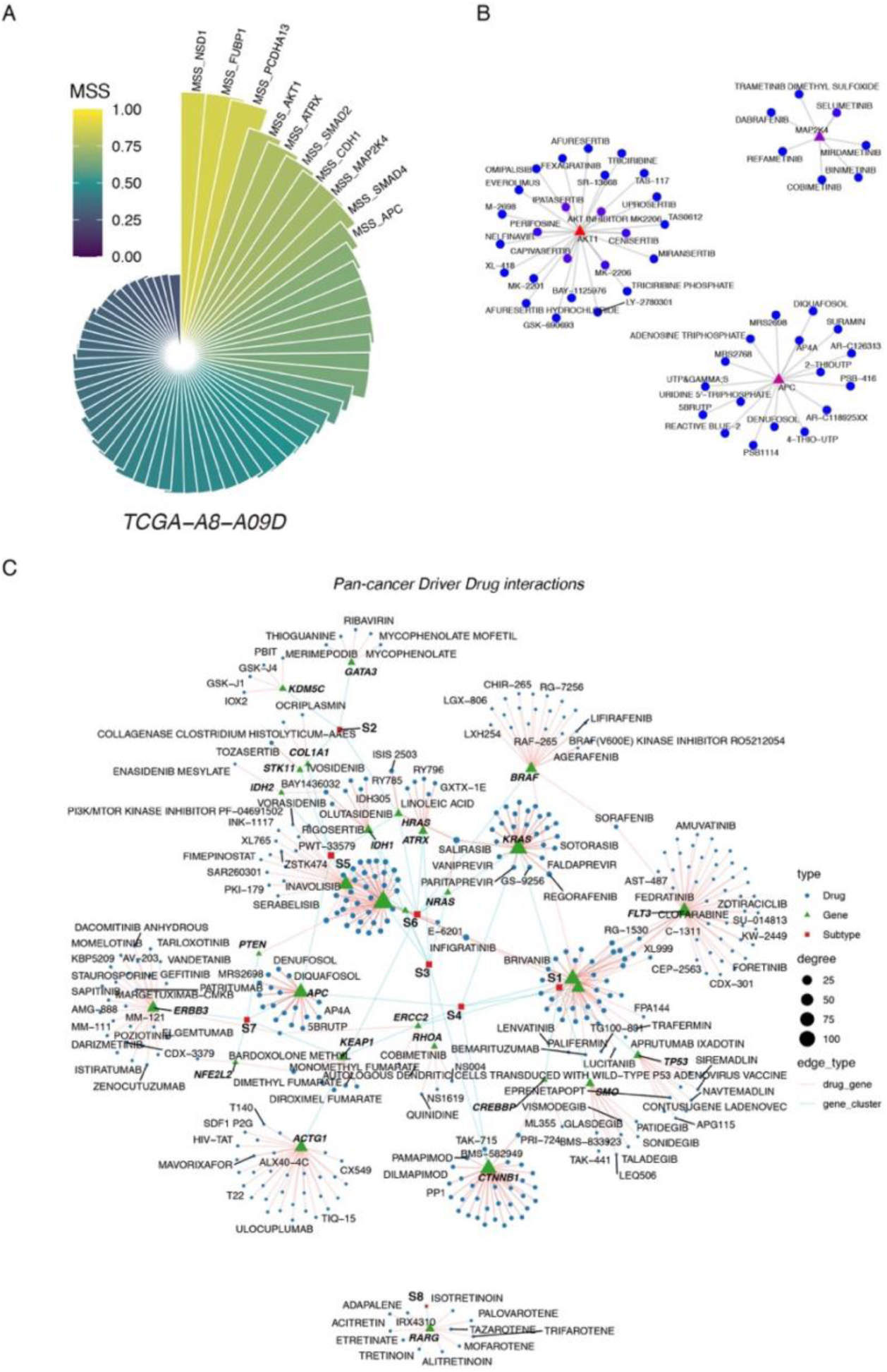
Driver mutation signature profiling enables patient-level interpretation and drug–target network mapping. (**A**) Radial bar plot showing mutation signature scores (MSS) for a representative tumor sample (TCGA-A8-A09D) across multiple driver genes. Each segment represents a driver-associated signature, colored by MSS value (scale 0–1). The plot highlights dominant driver-associated transcriptional programs within an individual tumor, illustrating the concept of a personalized mutation signature profile. (**B**) Network visualization of drug–target interactions for representative driver genes. Red triangles denote driver genes, and blue nodes represent drugs. Edges indicate known drug–gene interactions. (**C**) Global network integrating driver genes, therapeutic agents, and DMS subtypes. Green triangles represent driver genes, blue circles represent drugs, and red squares denote DMS subtypes (S1–S8). Edge types distinguish drug–gene interactions and gene–subtype associations. Node size corresponds to degree (number of connections). The network reveals subtype-specific therapeutic vulnerabilities and highlights central driver genes with extensive drug connectivity, supporting the potential of DMS-based stratification for guiding precision oncology.

Interestingly, the patient TCGA-A8-A09D is a breast cancer patient harboring a PIK3CA missense mutation (H1047R) (*33*), which explains an increased driver mutation signature of MAP2K4. This suggests that the patient might benefit from targeting PI3K/AKT signal transduction (**fig. 6B**, **fig. S10**).

By utilizing RNA-seq data in this way, precision cancer medicine can be enabled, as it allows for the extrapolation of specific driver mutations that are most relevant to an individual patient’s tumor, based on their expression data. This method can be used to predict the tumor’s molecular makeup and guide treatment strategies, even when somatic mutation data is not available. Furthermore, this approach can be applied broadly across diverse cancer types, enhancing personalized treatment strategies and improving patient outcomes (**fig. 6C**).

## Discussion

A central challenge in oncology is to define a molecular taxonomy of tumors that reflects their underlying biology rather than their tissue of origin (*34–40*). In this study, we show that driver mutation signatures (DMS)—the coordinated transcriptional programs induced by oncogenic drivers—provide such a framework. By systematically quantifying the downstream effects of driver mutations across cancers, we move beyond mutation-centric descriptions toward a systems-level representation of tumor identity. This approach reveals that tumors can be organized into discrete yet hierarchically structured subgroups, capturing both shared biological programs across lineages and divergence within individual cancer types.

Conceptually, our findings redefine the role of driver mutations: not merely as discrete genomic events (*41, 42*), but as architects of tumor state. Recurrent drivers such as TP53, KRAS and EGFR impose reproducible transcriptional programs that transcend tissue boundaries, enabling convergence of biologically similar tumors and separation of phenotypically distinct ones. This reconciles a long-standing paradox in tumor classification—why tumors of the same histology can behave differently, and why tumors from different tissues can share vulnerabilities—by anchoring classification in functional consequences rather than mutation presence alone.

Beyond classification, we propose that DMS represents a central integrative hub for multi-modal cancer data. Although derived from genomic (WES) and transcriptomic (RNA-seq) data (*43*), mutation signatures inherently encode the downstream consequences of upstream regulatory processes and the precursors of downstream molecular phenotypes. This concept is consistent with prior work showing that global transcription factor activity inferred from transcriptomic profiles can be used to stratify patients into clinically and biologically distinct subtypes with prognostic relevance(*44*). In this sense, DMS provides a natural interface to integrate additional data layers: epigenetic modifications that shape transcriptional outputs, proteomic states that execute cellular functions, and single-cell transcriptomic profiles that resolve cellular heterogeneity. Rather than viewing these modalities as independent dimensions, DMS offers a unifying axis along which they can be projected and interpreted, enabling a more coherent, cross-scale understanding of tumor biology.

The implications for precision oncology are substantial. As a biomarker, DMS subtyping extends beyond traditional mutation-based or histology-based classification by capturing the functional state of tumors. This enables patient stratification based on shared biological programs, potentially improving the prediction of therapeutic response. Importantly, DMS also facilitates the systematic mapping between drivers, downstream pathways, and drug sensitivities, providing a framework to mine driver–drug interactions at scale (*45–48*). In this context, mutation signature scores (MSS) can serve not only to group patients, but also to position individual tumors within a therapeutic landscape defined by pathway dependencies and actionable vulnerabilities.

At the same time, our results highlight an important principle: tumor subtyping is inherently hierarchical and, in principle, infinitely resolvable. While our analysis identifies robust, population-level subgroups, additional layers of heterogeneity exist within each class. Rather than treating subtypes as fixed categories, the continuous quantification of mutation signature scores provides a high-resolution fingerprint of each tumor. This “n = 1” representation captures the unique combination of driver-induced programs within an individual patient, offering a path toward truly personalized oncology where classification and treatment decisions are tailored at the single-tumor level.

Several limitations warrant consideration. Our framework is currently based on bulk transcriptomic data and thus reflects composite signals from tumor and microenvironmental compartments. Future integration with single-cell and spatial data will be essential to disentangle these contributions and refine the biological interpretation of mutation signatures. In addition, the robustness of signature definitions depends on cohort size and statistical thresholds, underscoring the need for standardized pipelines and prospective validation. Finally, clinical studies will be required to determine the predictive value of DMS-based stratification across diverse therapeutic settings.

In summary, we establish driver mutation signatures as a systems-level framework for tumor classification, bridging genotype and phenotype across cancers. By positioning DMS as both a unifying taxonomy and an integrative hub for multi-modal data, our study lays the foundation for a new generation of cancer biomarkers that operate across scales—from populations to individual patients—and supports a more precise and mechanistically grounded approach to cancer therapy.

## Methods

### Data Collection and Preprocessing

The data used in this study were sourced from The Cancer Genome Atlas (TCGA), a comprehensive resource containing genomic, transcriptomic, and clinical data from over 10000 tumor samples. RNA-seq expression data and mutation data were specifically chosen for this analysis. The RNA-seq data, provided as raw gene expression counts, were downloaded as .rds files (e.g., TCGA_<PROJECT>_RNA_STAR_Counts.rds), while mutation data were collected in Mutation Annotation Format (MAF) files, which contain detailed mutation information for each cancer sample. The mutation data were filtered to focus on driver genes, which are genes that, when mutated, contribute to tumorigenesis by driving essential biological processes like cell cycle regulation and apoptosis. The data were processed to remove low-quality genes with low counts across the samples and to harmonize gene identifiers by mapping Ensembl IDs to HGNC gene symbols using the biomaRt package (v2.52.0).

### Mutation Status Classification

For each cancer type, we classified the samples based on their mutation status in key driver genes. For each tumor sample, mutations in specific driver genes were identified, and the samples were labeled as either mutant or wildtype for each driver. This classification was based on the presence or absence of mutations in cancer driver genes. The mutation status for each sample was determined using the mutation data from the MAF files, where each sample’s mutation status in the driver genes was checked.

### Differential Expression Analysis

To evaluate the impact of each driver gene mutation on gene expression, we performed differential gene expression (DEG) analysis between the mutant and wildtype tumor samples using the DESeq2 (v1.36.0) (*49*). RNA-seq data for each cancer project were used to create a differentially expressed gene set, where samples were grouped based on their driver mutation status. The analysis aimed to identify genes that were significantly upregulated or downregulated in mutant samples compared to wildtype samples. The DEGs were selected based on log2 fold change (|log2FC| > 1) and adjusted p-value (padj < 0.05), indicating the genes most strongly associated with driver mutation status.

### Driver Mutation Signature Selection

The mutation signature for each driver gene was derived from the differentially expressed genes (DEGs) identified in the previous step. Genes that were upregulated in mutant samples across multiple projects were selected as part of the mutation signature for each driver gene. A minimum threshold was set for the number of projects in which a gene had to be upregulated to be included in the mutation signature. Only top upregulated genes (e.g., top 100) were retained, ensuring that only the most robust genes were used for the subsequent signature scoring. These mutation signatures were stored for each driver and used to calculate driver mutation signature scores.

### Driver Mutation Signature Scoring

Each tumor sample was assigned a mutation signature score based on the expression of the top mutation signature genes for each driver gene. This score was calculated by summing the expression levels of the upregulated signature genes for each sample. To ensure comparability across samples, the mutation signature scores were normalized to a 0–1 scale using min-max normalization. This approach provided a standardized way to evaluate each sample’s mutation signature profile relative to the entire cohort, enabling meaningful comparisons across cancer types.

### Clustering of Patients

Clustering of patients was performed based on their mutation signature scores, which reflected the presence of specific driver mutations in their tumor samples. The clustering was achieved using the k-means algorithm, where the number of clusters was determined using the elbow method, silhouette method, and gap statistic. These clustering methods were applied to the mutation signature scores in the Seurat object, allowing for the identification of distinct patient subgroups with shared mutation signature profiles. The number of clusters was selected based on the optimal balance between cluster cohesion and separation, as indicated by the silhouette score.

### Definition of driver mutation signature scores

To quantify the activity of driver mutation–associated transcriptional programs, we computed mutation signature scores (MSS) for each driver using predefined gene signatures derived from prior analyses. A union set of all signature genes across drivers was first constructed, and gene expression values were extracted from the RNA-seq matrix.

To ensure comparability across genes, gene-wise standardization was performed across all samples in the reference cohort. For each gene g, expression values were centered and scaled as:

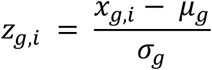

where *x_g,i_* denotes the expression of gene *g* in sample *i*, and *μ_g_* and *σ_g_* represent the mean and standard deviation of gene g across the reference cohort. Genes with zero variance were assigned a standard deviation of 1 to avoid numerical instability.

For each driver *d*, a raw mutation signature score was computed for each sample as the mean of standardized expression values across its signature gene set:

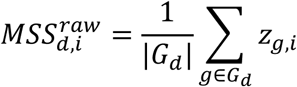

where *G_d_* denotes the set of genes associated with driver *d*. This score reflects the aggregate activation level of the mutation-associated transcriptional program.

To facilitate cross-sample comparison within the reference cohort, raw scores were transformed using a rank-based percentile normalization:

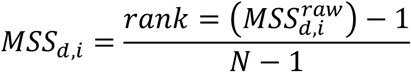

where *N* is the number of samples. This transformation yields values between 0 and 1, representing the relative activation of each driver signature across the cohort.

### Construction of the TCGA reference model

A pan-cancer reference model was constructed using bulk RNA-seq data from TCGA. For each driver signature, the following quantities were computed and stored: gene-level mean *μ_g_* and standard deviation *σ_g_* across TCGA samples; raw MSS scores for all samples; percentile-normalized MSS scores; driver-specific mean and standard deviation of raw MSS scores; subtype centroids defined as the average MSS profile within each MSS cluster; empirical cumulative distribution functions (ECDFs) of raw MSS scores for each driver. This reference model enables consistent projection of external datasets into the same MSS space.

### Pseudo-bulk construction from single-cell RNA-seq data

For scRNA-seq datasets, pseudo-bulk expression profiles were generated by aggregating raw counts across cells. Cells were grouped by sample or patient identifier, and gene-level counts were summed within each group:

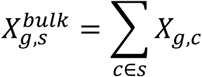

Where *X_g,c_* denotes the count of gene *g* in cell *c,* and s denotes a pseudo-bulk sample. Pseudo-bulk counts were then library-size normalized to counts per 10,000 and log-transformed:

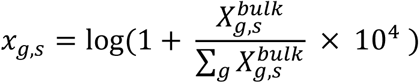

### Reference-anchored MSS scoring for query samples

To ensure comparability with the TCGA reference, MSS scores for query pseudo-bulk samples were computed using a reference-anchored approach. For each gene, expression values were standardized using the TCGA-derived parameters:

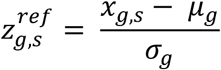

Driver-level raw scores were then computed as:

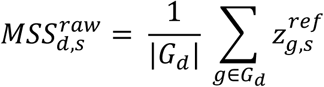

These raw scores were subsequently normalized using TCGA-derived distributions. Two transformations were used:

1. Z-score normalization relative to TCGA:

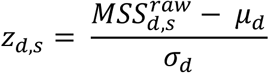

1. 2. Sigmoid transformation to bounded scale:

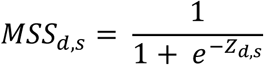

Alternatively, percentile scores were obtained by mapping raw scores to the empirical cumulative distribution of TCGA samples using ECDFs. This approach ensures that MSS values for query samples are directly comparable to the TCGA reference space.

### Subtype assignment by projection to reference MSS space

Subtype prediction for query samples was performed using two complementary strategies.

1. Centroid-based assignment

For each MSS subtype, a centroid was defined as the mean MSS profile across TCGA samples within that subtype. Query samples were assigned to subtypes based on Pearson correlation between their MSS profile and each centroid:

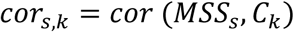

which denotes the centroid of subtype k. The subtype with the highest correlation was selected.

2. Nearest-neighbor assignment

Query samples were also compared to individual TCGA samples in MSS space using Euclidean distance. For each query sample, the k-nearest neighbors (k = 15) were identified, and subtype assignment was determined by majority vote among neighbors.

### Confidence assessment and interpretation

To evaluate the robustness of subtype assignments, we computed: Centroid margin: difference between the top and second-highest centroid correlations; kNN purity: proportion of nearest neighbors belonging to the assigned subtype.

Agreement between centroid-based and kNN-based predictions was used as an indicator of confidence. Discrepancies between methods were interpreted as evidence of intermediate or heterogeneous transcriptional states.

## Author contributions

Conceptualization: XH; Data curation: XH, BC, XQH; Methodology: XH, BC, XQH; Investigation: XH, BC, XQH; Visualization: XH, BC, XQH; Project administration: XH; Software: XH, BC, XQH; Validation: XH, BC, XQH; Supervision: XH, MW; Writing – original draft: XH; Writing – review & editing: XH, BC, XQH, MW.

## Data and materials availability

The transcriptomic and whole exome sequencing data from the Cancer Genome Atlas project were obtained from GDC data portal. The following datasets are available at GEO: GSE132465, GSE144735, GSE188711, GSE200997. The driver mutation signature scores generated in our study can be accessed at zenodo (DOI: 10.5281/zenodo.20264618) (*50*). Codes and intermediate results for the driver mutation signature pipeline is hosted at zenodo (DOI: 10.5281/zenodo.20280023) (*51*).

