## Supplementary Figures for "Pan-Cancer Driver Mutation Signatures Define a Molecular Taxonomy of Tumors"

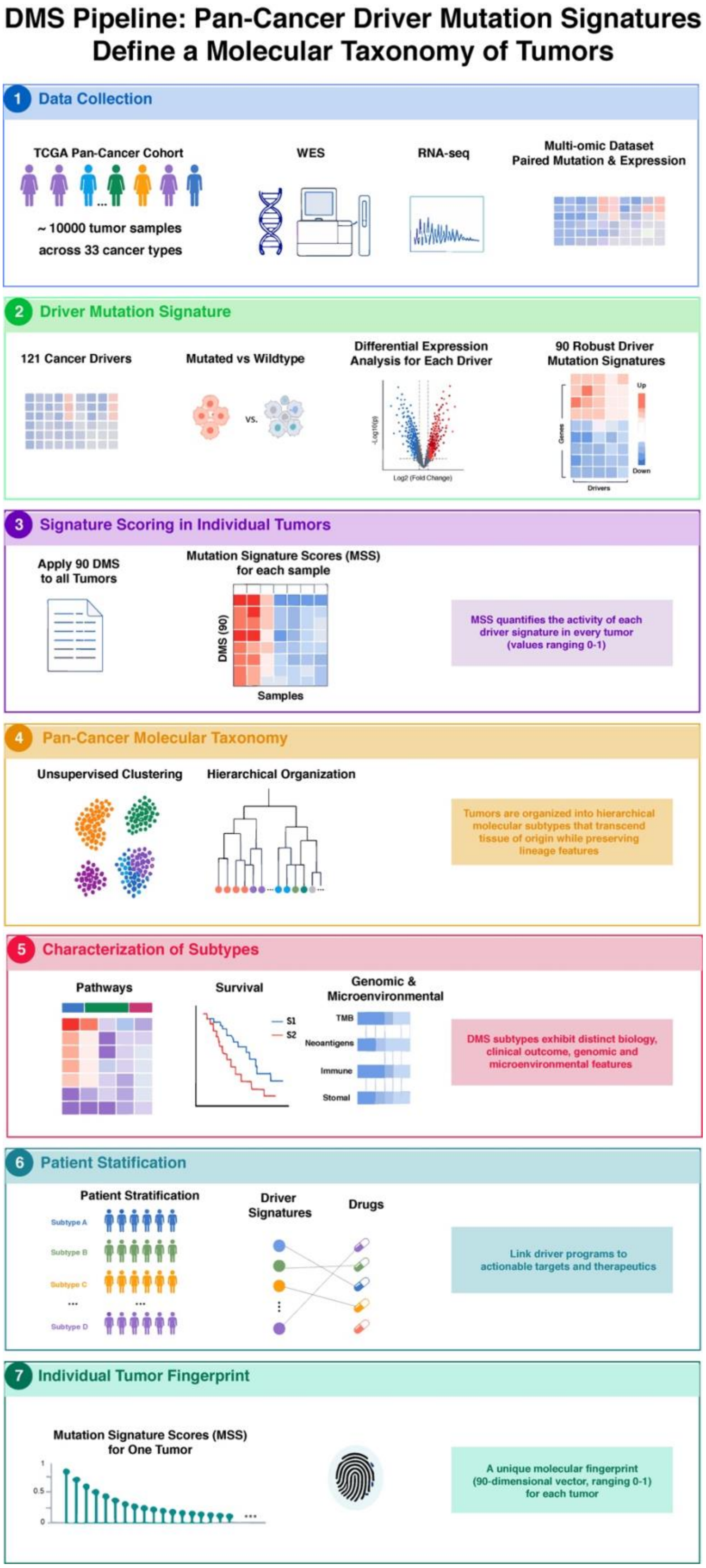
DMS Pipeline: Pan-Cancer Driver Mutation Signatures
Define a Molecular Taxonomy of Tumors
1 Data Collection
TCGA Pan-Cancer Cohort
~ 10000 tumor samples
across 33 cancer types
WES
RNA-seq
Multi-omic Dataset
Paired Mutation & Expression
2 Driver Mutation Signature
121 Cancer Drivers
Mutated vs Wildtype
vs.
Differential Expression
Analysis for Each Driver
-Log10(p)
Log2 (Fold Change)
90 Robust Driver
Mutation Signatures
Genes
Drivers
Up
Down
3 Signature Scoring in Individual Tumors
Apply 90 DMS
to all Tumors
Mutation Signature Scores (MSS)
for each sample
DMS (90)
Samples
MSS quantifies the activity of each
driver signature in every tumor
(values ranging 0-1)
4 Pan-Cancer Molecular Taxonomy
Unsupervised Clustering
Hierarchical Organization
Tumors are organized into hierarchical
molecular subtypes that transcend
tissue of origin while preserving
lineage features
5 Characterization of Subtypes
Pathways
Survival
S1
S2
Genomic &
Microenvironmental
TMB
Neoantigens
Immune
Stomal
DMS subtypes exhibit distinct biology,
clinical outcome, genomic and
microenvironmental features
6 Patient Statification
Patient Stratification
Subtype A
Subtype B
Subtype C
Subtype D
Driver
Signatures
Drugs
Link driver programs to
actionable targets and therapeutics
7 Individual Tumor Fingerprint
Mutation Signature Scores (MSS)
for One Tumor
1
0.5
0
A unique molecular fingerprint
(90-dimensional vector, ranging 0-1)
for each tumor

**Supplementary Figure S1. Overview of the DMS computational pipeline.**
Schematic illustration of the workflow for identifying pan-cancer driver mutation signatures (DMS) and deriving tumor molecular subtypes. The pipeline includes: (1) data collection from TCGA WES and RNA-seq datasets; (2) differential expression analysis for 121 pan-cancer drivers to define 90 robust DMS; (3) calculation of mutation signature scores (MSS) for each tumor; (4) unsupervised clustering and hierarchical organization to define DMS subtypes (S1–S8; S1.1-S1.8); (5) characterization of subtypes using pathway, immune, genomic, and clinical features; (6) patient stratification, linking driver signatures to potential therapeutic targets; and (7) derivation of individual tumor fingerprints as 90-dimensional MSS vectors. This workflow provides a systematic framework for integrating mutation-driven transcriptional programs with tumor heterogeneity, molecular subtype classification, and translational insights for precision oncology.

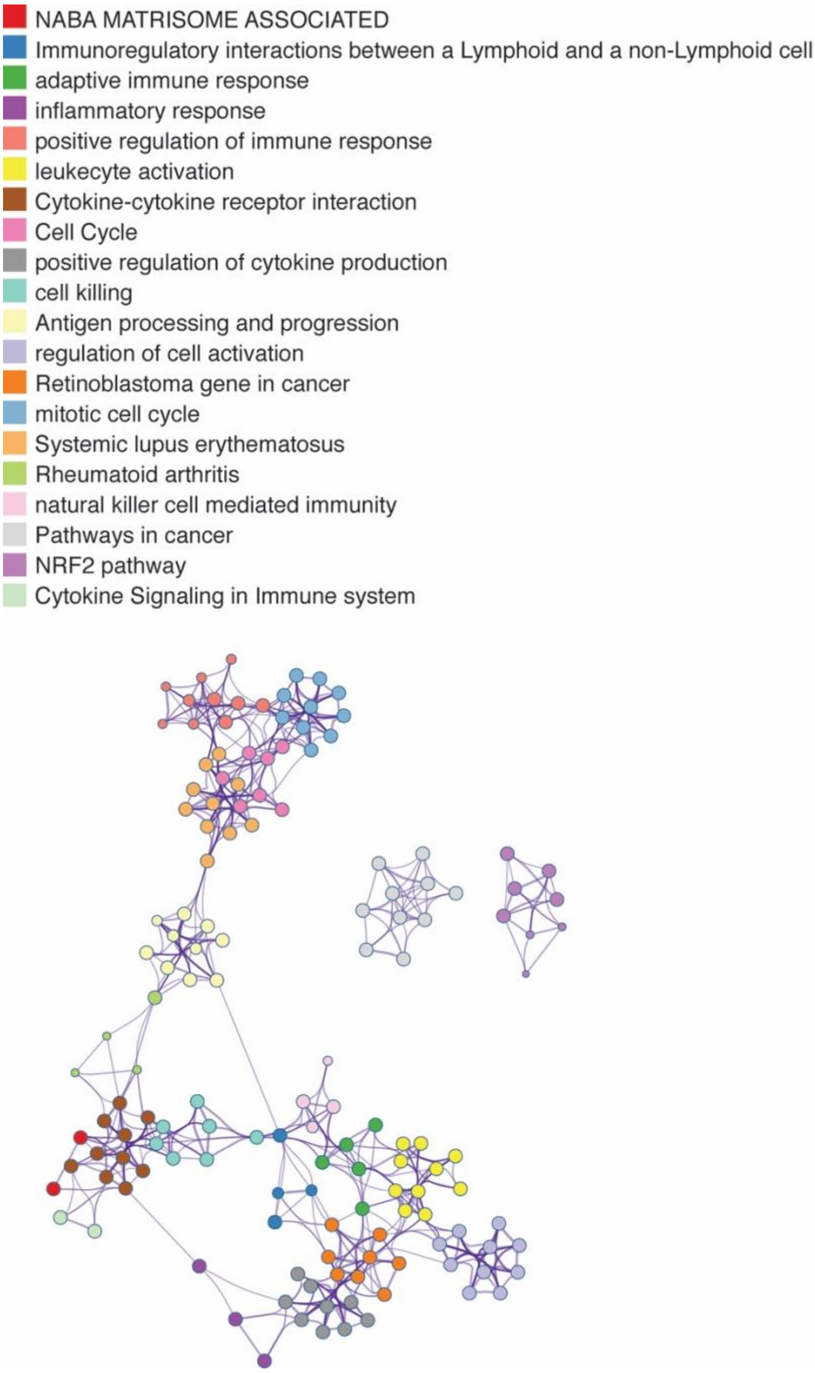
NABA MATRISOME ASSOCIATED
Immunoregulatory interactions between a Lymphoid and a non-Lymphoid cell
adaptive immune response
inflammatory response
positive regulation of immune response
leukecyte activation
Cytokine-cytokine receptor interaction
Cell Cycle
positive regulation of cytokine production
cell killing
Antigen processing and progression
regulation of cell activation
Retinoblastoma gene in cancer
mitotic cell cycle
Systemic lupus erythematosus
Rheumatoid arthritis
natural killer cell mediated immunity
Pathways in cancer
NRF2 pathway
Cytokine Signaling in Immune system

**Supplementary Figure S2. Functional enrichment network of driver mutation signatures.**
Network visualization of significantly enriched biological pathways associated with driver mutation signatures. Each node represents a pathway or gene set, colored according to functional category (e.g., immune response, cell cycle, cytokine signaling, extracellular matrix/matrisome). Edges indicate shared genes between pathways, reflecting functional overlap and coordinated biological programs.

Distinct clusters of pathways emerge, including immune-related processes (adaptive immune response, leukocyte activation, cytokine signaling), cell cycle and proliferation programs, and extracellular matrix organization (NABA matrisome). Additional modules highlight inflammation, antigen processing and presentation, and disease-associated pathways such as rheumatoid arthritis and systemic lupus erythematosus.

The network structure reveals that driver mutation signatures converge on a limited number of core biological processes, particularly immune regulation and proliferative signaling, supporting the functional coherence of DMS-derived transcriptional programs across cancers.

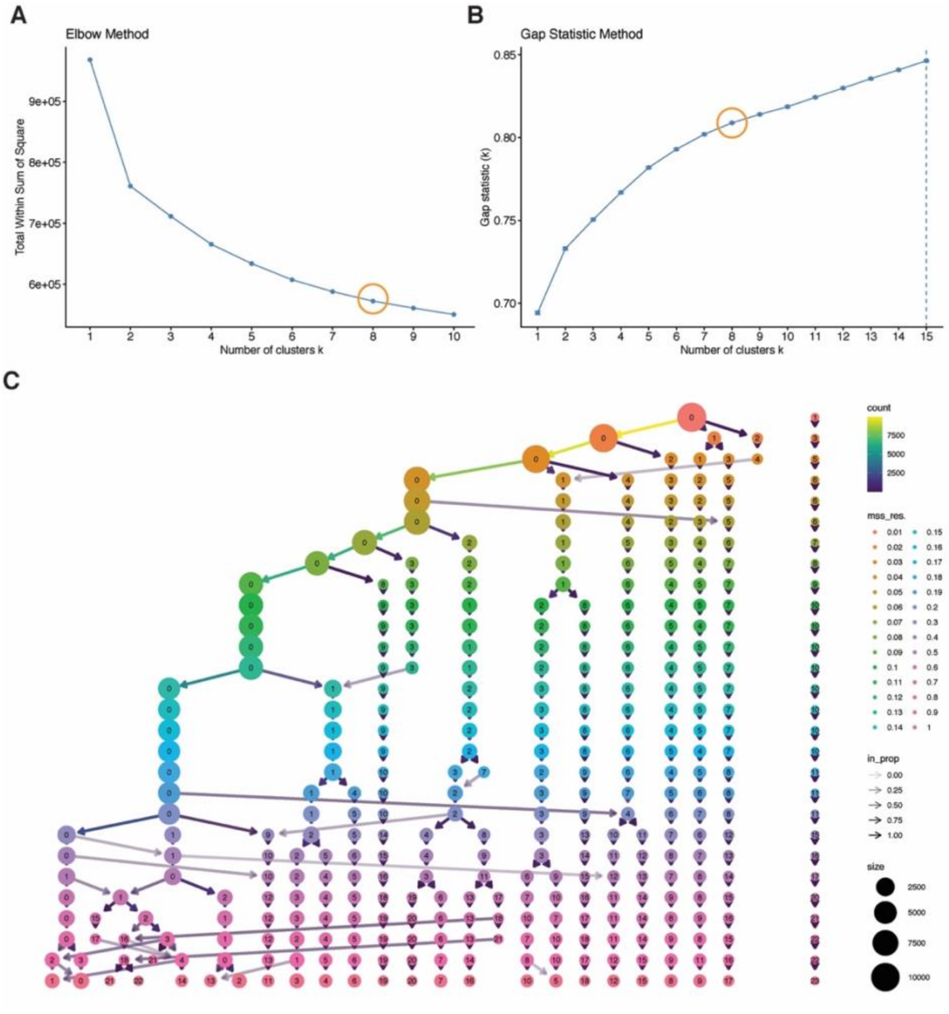


**Supplementary Figure S3. Determination and stability of DMS clustering.**
(A) Elbow method showing total within-cluster sum of squares across k, supporting k = 8. (B) Gap statistic analysis indicating an optimal clustering solution around k ≈ 8–9. (C) Cluster stability across resolutions visualized as a hierarchy, showing consistent mapping of samples and robustness of the DMS classification.

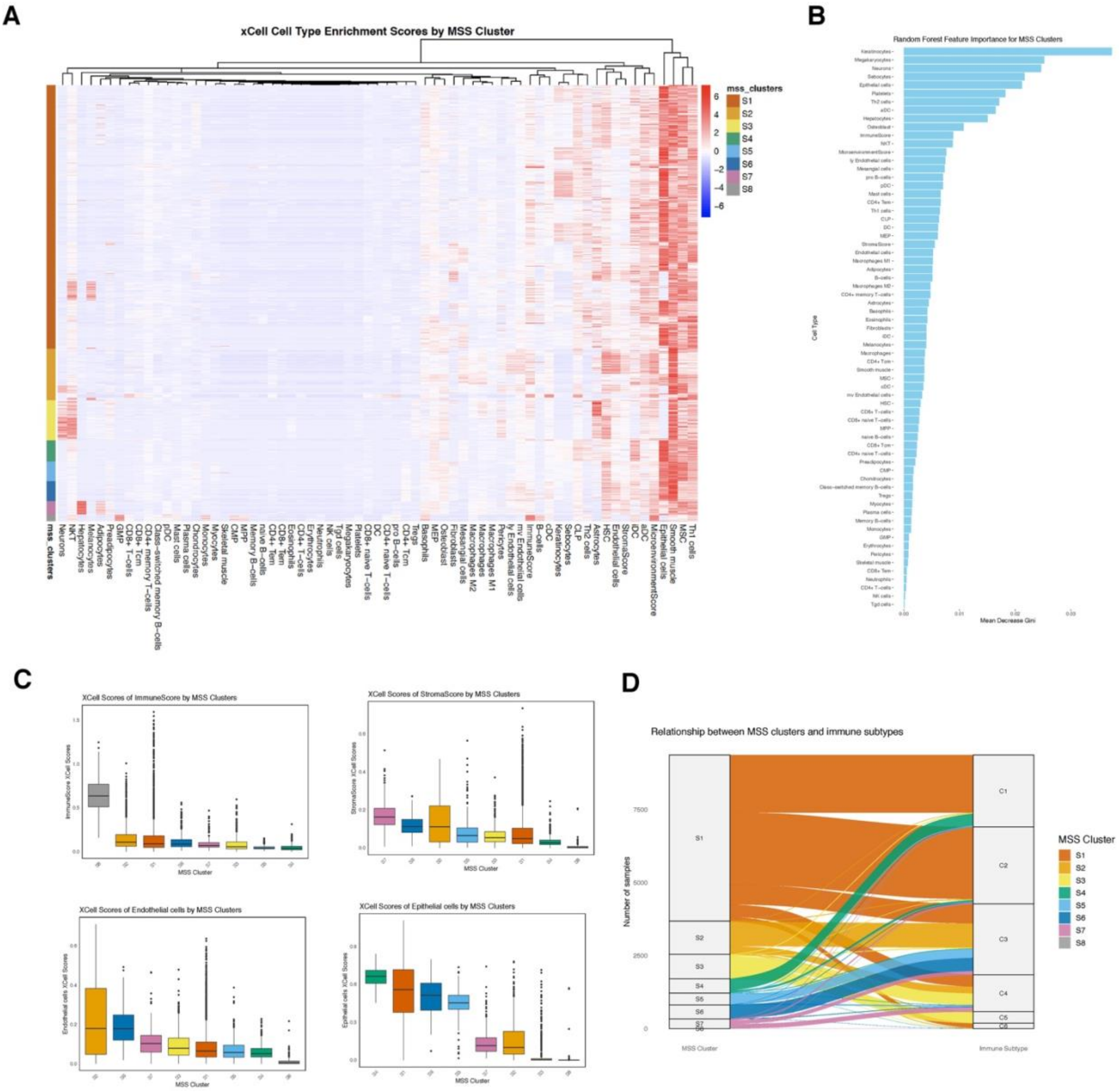


**Supplementary Figure S4. Immune and stromal features across DMS subtypes.**
(A) Heatmap of xCell-derived cell type enrichment scores across samples, grouped by DMS subtype (S1–S8). (B) Random forest feature importance ranking cell types contributing to DMS classification. (C) Distribution of immune, stromal, endothelial, and epithelial scores across subtypes. (D) Alluvial plot showing relationships between DMS subtypes and established immune subtypes (C1–C6).

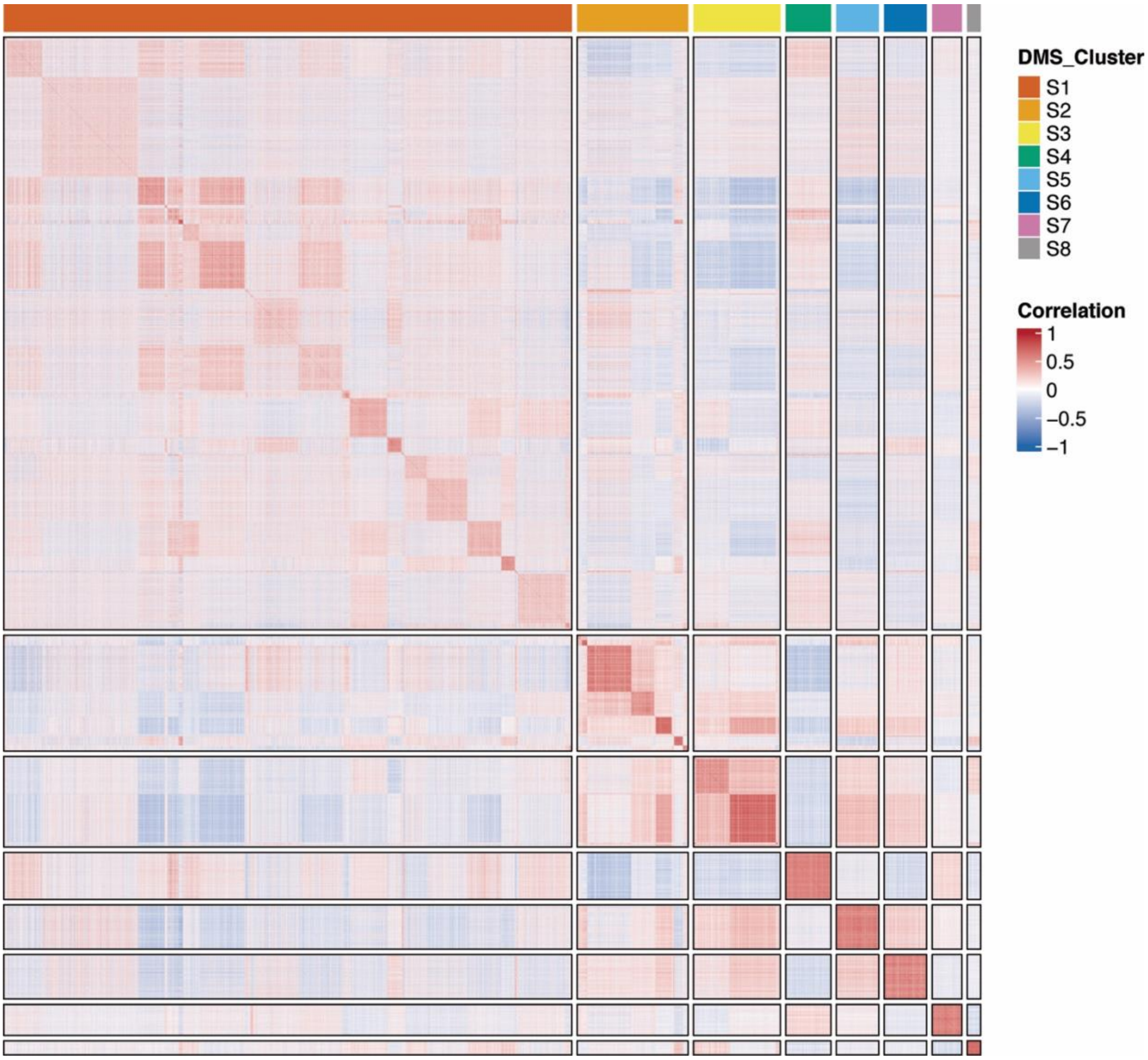


**Supplementary Figure S5. Sample–sample correlation of mutation signature scores.** Heatmap of pairwise correlations between tumor samples based on mutation signature scores (MSS). Samples are ordered by DMS subtype (S1–S8). Strong intra-subtype correlations and weaker inter-subtype correlations demonstrate the coherence and separability of DMS-defined molecular classes.

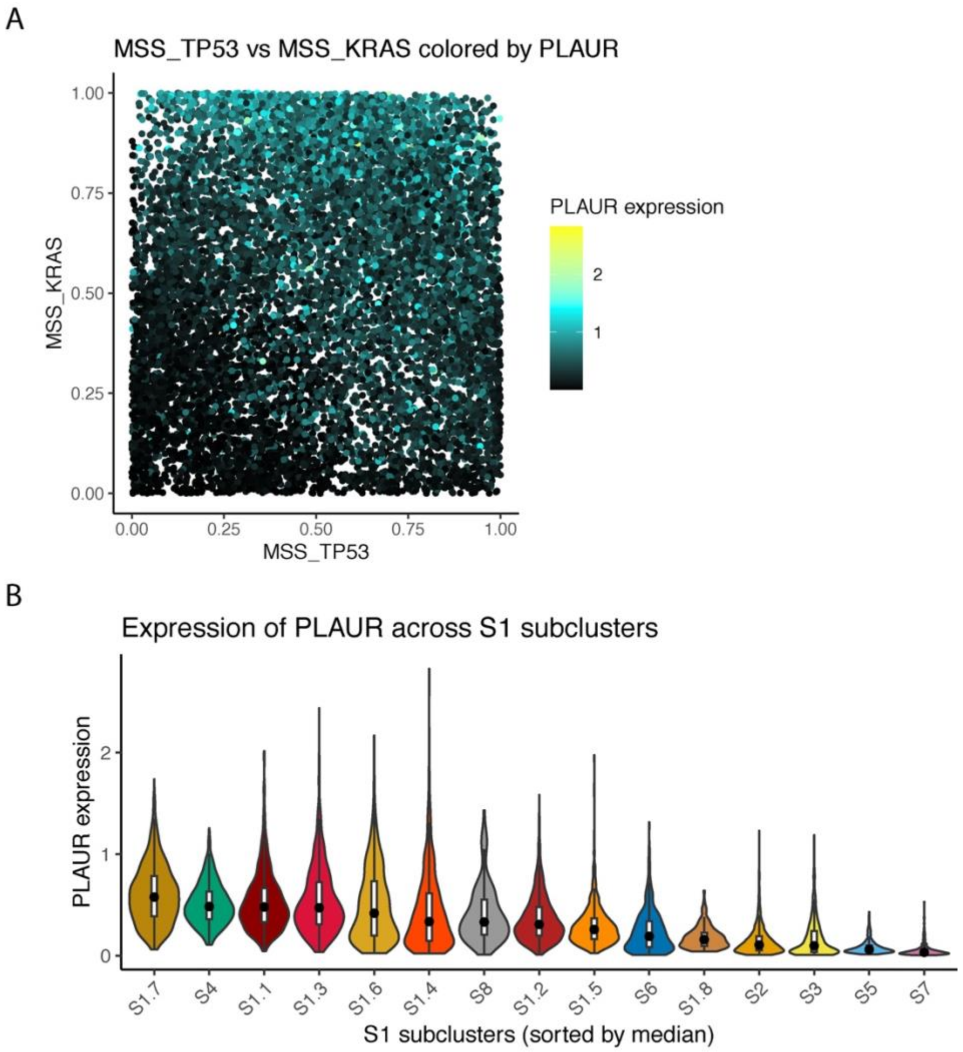


**Supplementary Figure S6. Expression of PLAUR in relation to TP53 and KRAS mutation signatures.** (A) Scatter plot of MSS_TP53 versus MSS_KRAS for pan-cancer tumor samples, with each point color-coded by PLAUR expression. High PLAUR expression is enriched in tumors with elevated TP53 and KRAS mutation signature activity. (B) Violin plot showing PLAUR expression across tumor subclusters, sorted by median.

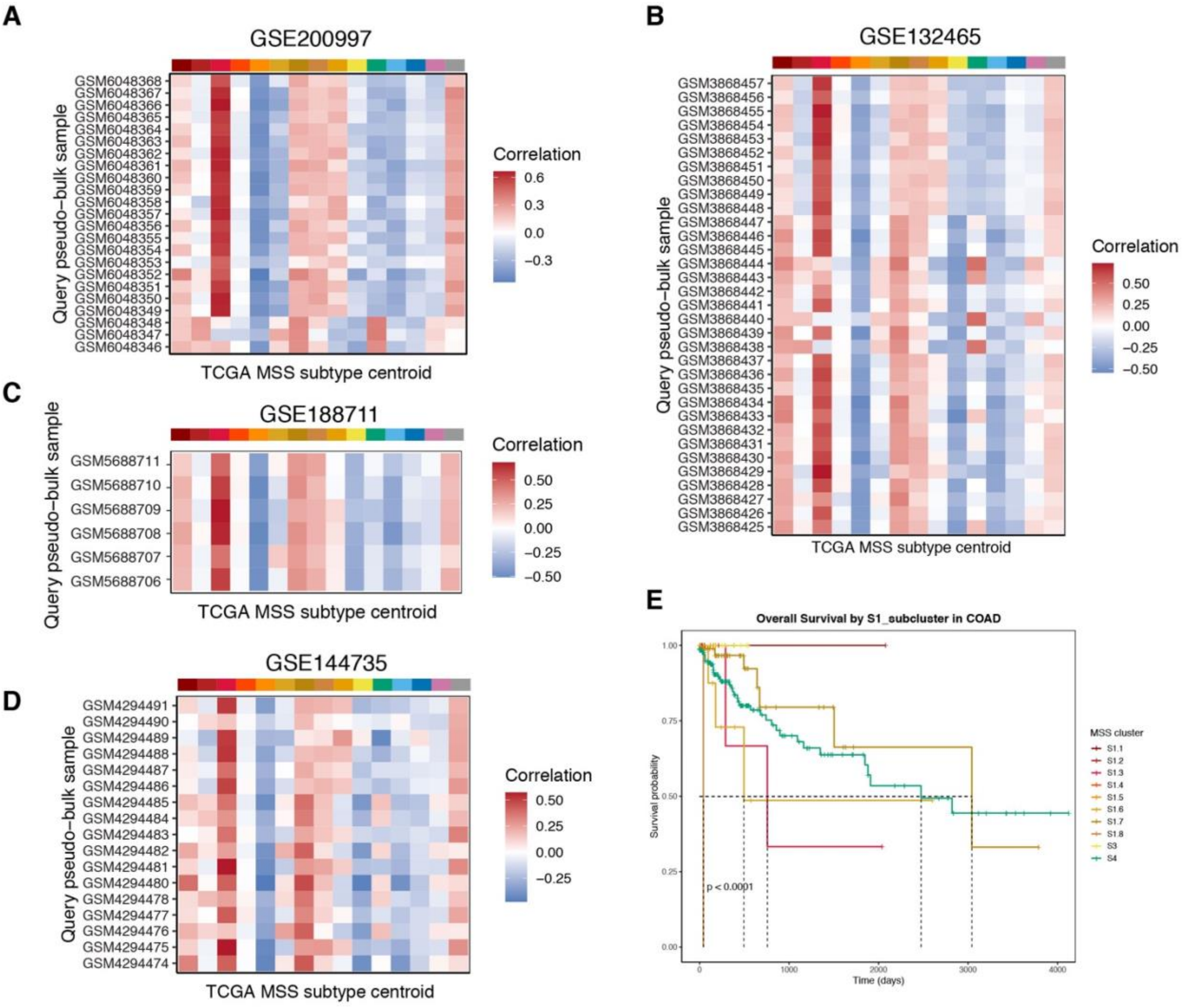


**Supplementary Figure S7. Validation of MSS-based subtyping using single cell RNA-seq datasets.** (A–D) Heatmaps showing the correlation between pseudo-bulk expression profiles of query samples and TCGA MSS subtype centroids for four independent datasets (GSE200997, GSE132465, GSE188711, GSE144735). Each row represents a pseudo-bulk query sample and each column represents the TCGA MSS subtype centroid. Color scale indicates Pearson correlation coefficients, with red representing positive correlation and blue negative correlation. (E) Kaplan–Meier analysis of overall survival in COAD patients based on tumor subcluster assignment. Survival curves are colored according to tumor subcluster identity.

| | Top Signatures | Druggable Hubs |
|---|---|---|
| **S1** | *TP53, NME8, CREBBP, FLT3, ZFP37, KRAS, FGFR3, ERCC2, RB1, SMO* | FGFR3 (eg. ERDAFITINIB, FUTIBATINIB, PAZOPANIB HYDROCHLORIDE); FLT3 (eg. GILTERITINIB, MIDOSTAURIN, QUIZARTINIB); KRAS (SOTORASIB, ADAGRASIB, SELUMETINIB) |
| **S2** | *GATA3, BAP1, HRAS, NSD1, BCOR, ZC3H13, COL21A1, KDM5C, STK11, PCDHA12* | GATA3 (eg. RIBAVIRIN, MYCOPHENOLATE, MERIMEPODIB); KDM5C (IOX2, GSK-J1, PBIT); STK11 (TOZASERTIB) |
| **S3** | *IDH1, BCOR, NRAS, MAP3K1, PIK3R1, ATRX, BAP1, RHOA, CTNNB1, SMAD2* | PIK3R1 (eg. PANULISIB, INFIGRATINIB, RECILISIB); CTNNB1 (eg. AZD7624, BMS-582949, LOSMAPIMOD); IDH1 (eg. VORASIDENIB, IDH305, IVOSIDENIB) |
| **S4** | *APC, FGFR2, KRAS, CTNNB1, KEAP1, FGFR3, FBXW7, SETD2, ABCD3, ZNF624* | FGFR2/FGFR3 (eg. ERDAFITINIB, FUTIBATINIB, PAZOPANIB HYDROCHLORIDE); CTNNB1 (eg. AZD7624, BMS-582949, LOSMAPIMOD); KRAS (SOTORASIB, ADAGRASIB, SELUMETINIB) |
| **S5** | *COL1A1, PIK3R1, MAP3K1, TXNIP, LZTR1, IDH2, APC, PIK3CA, HRAS, STK11* | PIK3R1/PIK3CA (eg. RIGOSERTIB, ALPELISIB, SONOLISIB); APC (AP4A, MRS2698, ADENOSINE TRIPHOSPHATE) |
| **S6** | *HRAS, IDH1, PIK3R1, ATRX, ACTG1, NRAS, CDKN2A, SF3B1, BRAF, KRAS* | PIK3R1 (eg. PANULISIB, INFIGRATINIB, RECILISIB); ACTG1 (eg. MOTIXAFORTIDE, PLERIXAFOR); BRAF (eg. DABRAFENIB, BELVARAFENIB, DASATINIB); KRAS (eg. SOTORASIB, ADAGRASIB, SELUMETINIB) |
| **S7** | *KEAP1, SMAD2, NFE2L2, ABCD3, APC, SETD2, PCDHA12, PTEN, ERBB3, CTCF* | ERBB3 (eg. SERIBANTUMAB, PATRITUMAB, ZENOCUTUZUMAB); APC (AP4A, MRS2698, ADENOSINE TRIPHOSPHATE); KEAP1/NFE2L2 (BARDOXOLONE METHYL) |
| **S8** | *SF3B1, CDKN1A, ZC3H13, RARG, TBX3, ZNF750, RAC1, CDH1, RUNX1, FCRL4* | RARG (eg. TAZAROTENE, TRIFAROTENE, MOFAROTENE) |

**Supplementary Figure S8. Summary of subtype-specific driver signatures and druggable targets.**Table summarizing key driver mutation signatures and representative druggable hubs across DMS subtypes (S1–S8). For each subtype, top contributing driver genes and corresponding therapeutic targets are listed, highlighting subtype-specific molecular features and potential treatment strategies.

| | Top Signatures | Druggable Hubs |
|---|---|---|
| **S1.1** | *FGFR3, CDKN2A, FLT3, NOTCH1, STAG2, ABCD3, FBXW7, KDM6A, SMO, TAS1R3* | FGFR3 (eg. ERDAFITINIB, FUTIBATINIB, PAZOPANIB HYDROCHLORIDE); FLT3 (eg. GILTERITINIB, MIDOSTAURIN, QUIZARTINIB); SMO (eg. SONIDEGIB, TALADEGIB, VISMODEGIB); NOTCH1 (eg. BRONTICTUZUMAB, ENOTICUMAB) |
| **S1.2** | *AJUBA, PDYN, MAP2K1, KLF5, RXRA, ACVR1B, ZNF624, PTEN, CEACAM8, CTCF* | MAP2K1 (eg. SELUMETINIB, TRAMETINIB DIMETHYL SULFOXIDE, MIRDAMETNIB); RARA (eg. ETRETINATE, MOFAROTENE, ACITRETIN); |
| **S1.3** | *EP300, CASP8, MTOR, CDH1, BRCA1, PCDHA12, ERBB3, EGFR, SMO, TAS1R3* | EGFR (eg. NIMOTUZUMAB, OSIMERTINIB, AFATINIB); ERBB3 (eg.ISTIRATUMAB, ZENOCUTUZUMAB, ELGEMTUMAB); MTOR (eg. METFORMIN, OMIPALISIB, DACTOLISIB) |
| **S1.4** | *KLF5, NME8, SETD2, TP53, RUNX1, ACTG1, FOXA1, RARG, CREBBP, TAS1R3* | ACTG1 (eg. MOTIXAFORTIDE, PLERIXAFOR); TP53 (eg. NAVTEMADLIN, GRANISETRON) |
| **S1.5** | *AKT1, MAP2K4, RHOA, PIK3R1, TXNIP, ACTG1, KRAS, COL1A1, ERCC2, KLF5* | PIK3R1 (eg. PANULISIB, INFIGRATINIB, RECILISIB); AKT1 (eg. CAPIVASERTIB, IPATASERTIB, TAS0612); KRAS (eg. SOTORASIB, ADAGRASIB, SELUMETINIB); ACTG1 (eg. MOTIXAFORTIDE, PLERIXAFOR) |
| **S1.6** | *KRAS, FGFR3, WT1, STAG2, CDKN2A, ERBB2, KEAP1, FGFR2, FBXW7, TXNIP* | FGFR2/FGFR3 (eg. ERDAFITINIB, FUTIBATINIB, PAZOPANIB HYDROCHLORIDE); KRAS (eg. SOTORASIB, ADAGRASIB, SELUMETINIB); ERBB2 (eg. LAPATINIB, RITUXIMAB, PYROTINIB) |
| **S1.7** | *KRAS, FGFR2, LZTR1, RXRA, RAC1, PDYN, FCRL4, CEACAM8, NOTCH1, MAP2K1* | MAP2K1 (eg. SELUMETINIB, TRAMETINIB DIMETHYL SULFOXIDE, MIRDAMETNIB); FGFR2 (eg. LENVATINIB, INFIGRATINIB, ERDAFITINIB); KRAS (eg. SOTORASIB, ADAGRASIB, SELUMETINIB) |
| **S1.8** | *TP53, NME8, NRAS, NSD1, ERCC2, RUNX1, ZFP37, FBXW7, CIC, CREBBP* | TP53 (eg. NAVTEMADLIN, GRANISETRON) |

**Supplementary Figure S9. Subcluster-specific driver signatures and druggable targets within S1.** Table summarizing top driver mutation signatures and representative druggable hubs across S1 subclusters (S1.1–S1.8). Distinct combinations of driver genes and therapeutic targets highlight refined molecular heterogeneity and potential subtype-specific treatment opportunities within S1.

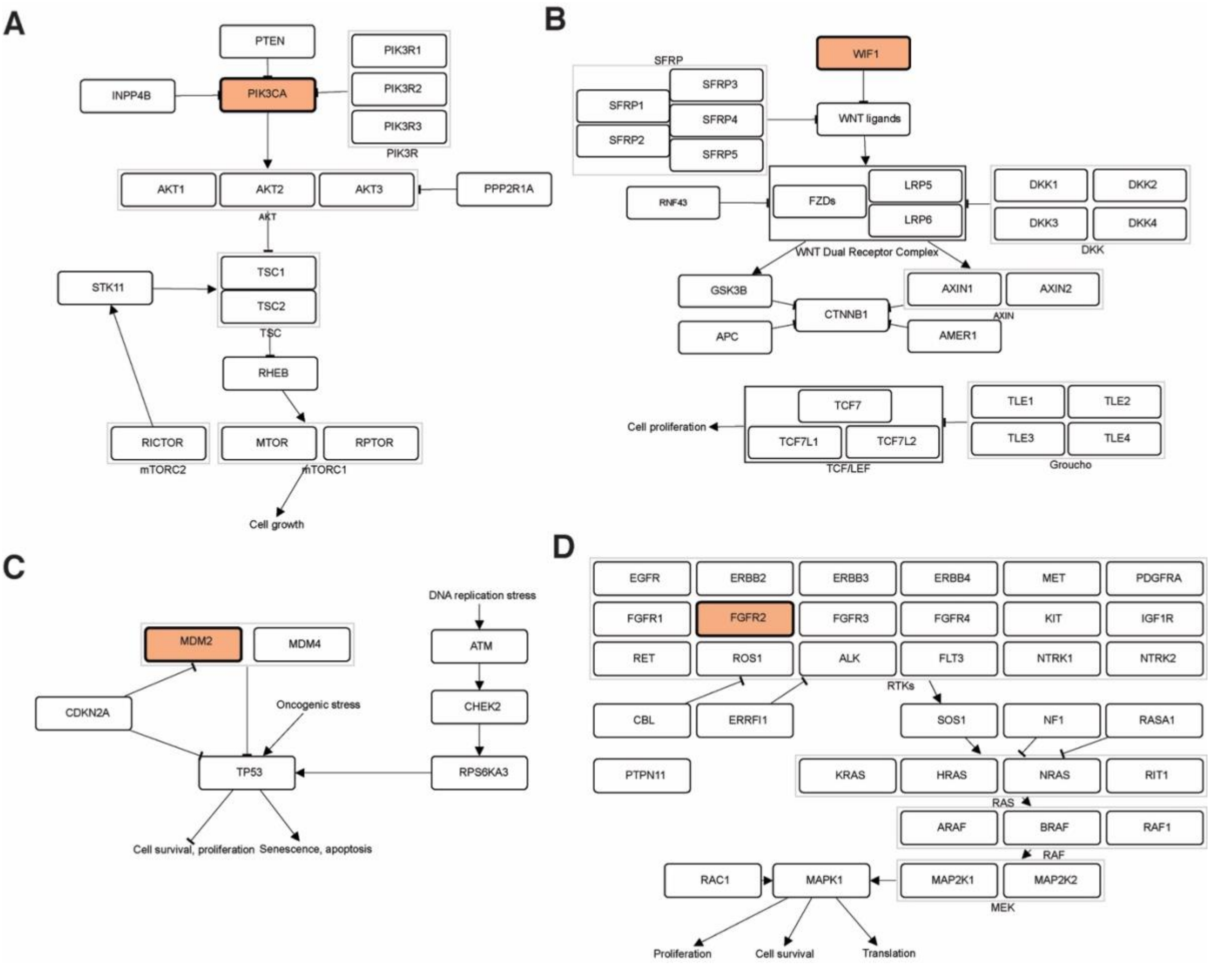


**Supplementary Figure S10. Mutated signaling pathways in a representative patient (TCGA-A8-A09D).** Pathway diagrams highlighting somatic mutations identified by whole-exome sequencing in key oncogenic pathways: (A) PI3K–AKT–mTOR, (B) WNT/β-catenin, (C) p53, and (D) RTK–RAS–MAPK. Mutated genes are highlighted, illustrating patient-specific alterations across major cancer signaling networks.
